# Targeted replacement of full-length CFTR in human airway stem cells by CRISPR/Cas9 for pan-mutation correction in the endogenous locus

**DOI:** 10.1101/2021.02.26.432961

**Authors:** Sriram Vaidyanathan, Ron Baik, Lu Chen, Dawn T. Bravo, Carlos J. Suarez, Shayda M. Abazari, Ameen A. Salahudeen, Amanda M. Dudek, Christopher A. Teran, Timothy H. Davis, Ciaran M. Lee, Gang Bao, Scott H. Randell, Steven E. Artandi, Jeffrey J. Wine, Calvin J. Kuo, Tushar J. Desai, Jayakar V. Nayak, Zachary M. Sellers, Matthew H. Porteus

## Abstract

Cystic fibrosis (CF) is a monogenic disease caused by impaired production and/or function of the cystic fibrosis transmembrane conductance regulator (CFTR) protein. Although we have previously shown correction of the most common pathogenic mutation, there are many other pathogenic mutations throughout the CF gene. An autologous airway stem cell therapy in which the CFTR cDNA is precisely inserted into the CFTR locus may enable the development of a durable cure for almost all CF patients, irrespective of the causal mutation. Here, we use CRISPR/Cas9 and two adeno-associated viruses (AAV) carrying the two halves of the CFTR cDNA to sequentially insert the full CFTR cDNA along with a truncated CD19 (tCD19) enrichment tag in upper airway basal stem cells (UABCs) and human bronchial basal stem cells (HBECs). The modified cells were enriched to obtain 60-80% tCD19^+^ UABCs and HBECs from 11 different CF donors with a variety of mutations. Differentiated epithelial monolayers cultured at air-liquid interface showed restored CFTR function that was >70% of the CFTR function in non-CF controls. Thus, our study enables the development of a therapy for almost all CF patients, including patients who cannot be treated using recently approved modulator therapies.

## Introduction

Cystic fibrosis (CF), one of the most common life-shortening genetic diseases, can result from a variety of deleterious mutations spanning the cystic fibrosis transmembrane conductance regulator (CFTR) gene. Among the >1,700 known CFTR mutations that can cause CF, F508del affects most CF patients in Europe and North America. Small molecule CFTR modulators that aid CFTR folding or potentiation increase CFTR function and improve health outcomes in patients with at least one F508del allele or certain gating mutations, comprising ∼90% of the known CF population ^1–3^. However, significant interpatient heterogeneity of response has been observed in these modulators^1–3^. In addition, modulators must be administered daily, are costly, only partially restore function, and do not cover the full spectrum of CFTR mutations ^2,4,5^. Thus, there is a continued need to develop a durable therapy that can treat 100% of CF patients.

Several gene therapy trials for CF have been performed ^6–8^. Although CF is a systemic disease, past attempts to develop a gene therapy have focused on the respiratory tract because of the high morbidity and mortality associated with CF lung disease. These trials used viral or non-viral vectors to deliver a copy of the CFTR cDNA into the cells of the epithelia in the upper (nose, sinuses) and lower airways ^6–8^. These trials were unsuccessful in producing a durable improvement in pulmonary function in CF patients. Although in vivo genome editing is being attempted to treat other diseases (e.g. muscular dystrophy),^9^ immune responses against gene editing enzymes such as Cas9 have been shown to limit the persistence of gene corrected cells in mouse models ^10^. We pursued the development of an autologous gene corrected airway stem cell therapy to treat CF in order to avoid anti-Cas9 adaptive immune responses which have been shown to exist in humans _11,12_.

A major obstacle to the successful development of ex vivo gene therapy is the inability to gene correct CFTR mutations or add a functional copy of the CFTR cDNA safely with high efficiencies in airway basal stem cells. Several prior studies have attempted to insert the CFTR cDNA in airway basal stem cells and airway tissue using integrating viruses ^13,14^. In these studies, the expression of CFTR was driven by non-native promoters and the integration of CFTR in the genome occurred in a semi-random manner. Nevertheless, these studies showed that the overexpression of CFTR using non-native promoters was sufficient to restore CFTR function, both in vitro and in vivo. In this study, we show the precise insertion of the CFTR cDNA in the endogenous CFTR locus, thereby preserving natural CFTR expression.

The discovery and development of zinc finger nucleases, transcription activator-like effective nucleases (TALEN) and Cas9 have enabled the precise insertion of exogenous DNA sequences into the genome ^15,16^. The ex vivo gene correction of F508del and other CFTR mutations using nuclease mediated editing has been performed in induced pluripotent stem cells (iPSC) and primary intestinal stem cells ^17–19^. Although the differentiation of iPSCs into CFTR-expressing airway epithelia consisting of cells resembling ciliated, secretory and basal cells has been shown ^20,21^, long-term differentiation potential of the iPSC derived basal cells still needs to be validated ^22^. As an alternative, we recently reported an ex vivo cell therapy approach using airway stem cells derived from the nasal and sinus epithelia, termed upper airway basal stem cells (UABCs) as well as human bronchial epithelial cells (HBECs) ^23^. Using Cas9 and adeno-associated virus (AAV), we achieved >40% correction of the F508del mutation in a selection-free manner and observed 30-50% restoration of CFTR function relative to non-CF controls by Ussing chamber analysis ^23^. This approach can be adapted to correct other mutations individually. However, because numerous non-F508del mutations, each relatively rare, are responsible for CF in patients who are not helped by CFTR modulators, a more efficient gene-therapy strategy might be the insertion of the entire CFTR cDNA into the endogenous locus.

One challenge in using Cas9/AAV to insert the CFTR cDNA is that the CFTR cDNA (4500 bp) is close to the packaging limit of AAV (4800 bp). As a result, there is insufficient space for the inclusion of homology arms and other components such as a poly-A sequence or a detection tag to enrich for the edited basal stem cells. To overcome this challenge, we applied a previously reported strategy in which sequences >5000 bp in length were inserted sequentially using two templates containing two halves of the HR template ^24^. In this study, we use this strategy to insert the CFTR cDNA and an enrichment cassette expressing truncated CD19 (tCD19) in the endogenous CFTR locus of airway basal stem cells ^24^. Thus, our strategy enables the development of a universal gene therapy (Universal strategy) for virtually all CF patients and addresses a significant hurdle in the development of ex vivo gene therapies for CF.

## Results

### Optimized expansion of primary upper airway basal cells

We previously reported the expansion of UABCs as monolayers in media containing epidermal growth factor (EGF), the bone morphogenetic protein (BMP) antagonist Noggin, and the transforming growth factor-β (TGF-β) inhibitor A83-01 (EN media) in plates coated with tissue culture collagen/laminin basement membrane extract (BME) ^23^. However, BME cannot be used to produce clinical grade cell therapy products because it is derived from mouse sarcoma cells. Hence, we sought to identify an alternative coating material that is clinically compatible.

Upper airway tissues were obtained from CF and non-CF patients undergoing functional endoscopic sinus surgeries (FESS). UABCs were isolated by digestion with pronase as previously reported and expanded using EN media ^23^. We screened the use of amine, carboxyl, fibronectin functionalized plates (data not shown) and a recombinant laminin 511 coating (iMatrix™). The use of iMatrix™ coating improved the expansion of UABCs by a factor of 1.5 ± 0.4 with respect to UABCs cultured in plates coated with BME extract (Figure 1A). To test the impact of iMatrix™ coating on the expansion of genome edited UABCs, we edited non-CF UABCs using our previously reported reagents specific to the F508del mutation and expanded the edited UABCs in plates coated with BME or iMatrix™ ^23^. Edited UABCs cultured in iMatrix™ coated plates showed improved expansion consistent with our observations in experiments with unmanipulated UABCs (Figure 1B). All further experiments used UABCs plated on tissue culture plates coated with iMatrix™.

**Fig. 1.**
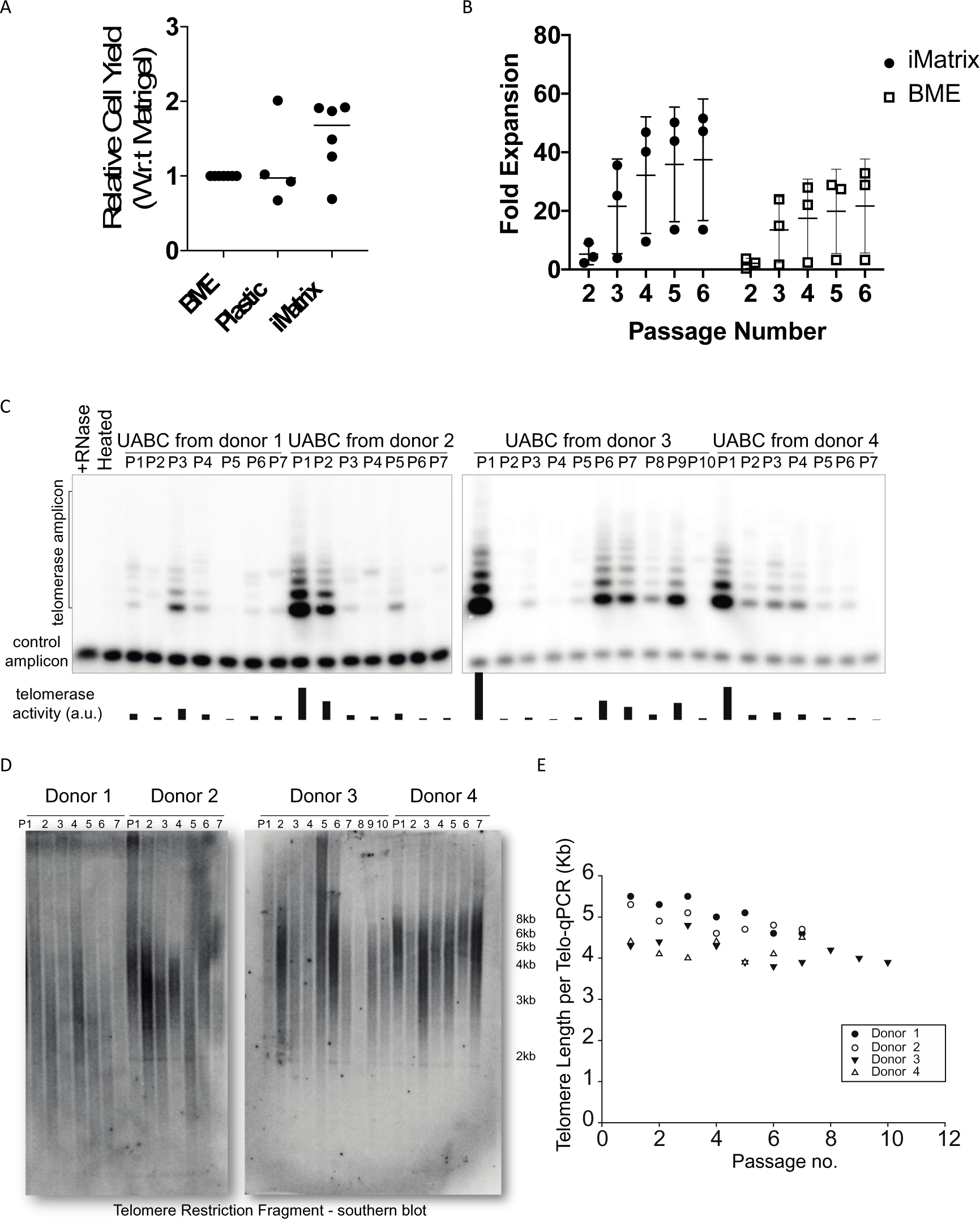
Expansion and characterization of UABCs. (**A**) UABCs cultured in tissue cultured plates coated with iMatrix™ (recombinant laminin 511) showed improved proliferation immediately after plating from tissue and (**B**) during subsequent passages. (**C**) Telomerase activity in the Expanded UABCs was measured using Telomeric Repeat Amplification Protocol (TRAP) and telomerase activity was observed in UABCs across multiple passages. (**D**) Significant shortening of telomere lengths was also not observed in expanded UABCs when telomere lengths were probed using Southern blot or (**E**) qPCR

Since UABCs expanded significantly on iMatrix, we were concerned about the possibility of extensive telomere erosion which could be detrimental to the long-term durability of a cell therapy. Hence, we measured telomerase expression and telomere lengths in UABCs cultured ex vivo. We observed telomerase activity in UABCs across multiple passages spanning 3 weeks in multiple donors (Figure 1C) and we did not observe a significant shortening of telomere lengths as measured by Southern blot and qPCR (Figures 1D-E).

### Optimizing the insertion of the full-length CFTR cDNA and enrichment of corrected basal stem cells

We first evaluated the possibility of inserting the CFTR cDNA in the endogenous locus using a single AAV. Genome editing using Cas9 results in double stranded breaks (DSBs) that are repaired either using non-homologous end joining (NHEJ) or homologous recombination (HR). The insertion of the CFTR cDNA requires the HR pathway which relies on correction templates containing the cDNA of interest flanked by homology arms (HA) that resemble the DSB site. Previous studies reporting high levels of HR in primary cells using AAV reported HAs of ≥400 bp ^25–27^. Because the CFTR cDNA (∼4500 bp) is close to the packaging capacity of AAV (4800-4900 bp), there is insufficient room to include HAs of 400 bp. However, CFTR can be packaged into AAV with small homology arms of ∼50 bp and a short polyA sequence (<100 bp).

To determine if efficient HR can be achieved with small HAs, we used HR templates coding for green fluorescent protein (GFP), but not CFTR, containing HAs of 50, 100, 150 and 400 bp. This GFP expression cassette utilized the spleen focus forming virus (SFFV) promoter. In UABCs edited using these GFP expressing templates, we observed that only 10% of UABCs treated with templates containing HA of 50 bp were GFP^+^ and only 15% GFP^+^ UABCs were produced when using HR templates with 100 bp HAs (Figure S1A). Since previous studies showed that allelic correction levels above 20% were most consistent in restoring CFTR function ^23^, we pursued an alternate strategy that would enable the enrichment of corrected UABCs.

To enable the inclusion of longer HAs and additional genetic sequences that allow the enrichment of corrected cells, we adapted a previously reported strategy in which two halves of the full template are packaged into two separate AAV vectors which then undergo fusion at the target genomic site by HR following a DSB ^24^. In this strategy, two sequential HR events as illustrated in Figure S2 will result in the insertion of the CFTR cDNA and an enrichment cassette containing tCD19 under the control of the PGK promoter (Figure 2A). The use of truncated human cell surface proteins such as CD19 and the nerve growth factor receptor (tNGFR) to enrich genetically modified cells for clinical applications has been previously reported ^28–32^. We used tCD19 in this study because airway basal stem cells are known to endogenously express NGFR. Notably, the tCD19 protein does not contain the intracellular signaling domains present in the endogenous protein and not capable of affecting cell phenotype by intracellular signaling. We do note that the extracellular portion of CD19 could be a target for CAR-T therapies (Yescarta and Kymriah) or bi-specific antibody therapy (blinatumomab) if it ever became necessary to eliminate the engineered cells and their progeny as a possible safety feature. Lastly, we used the PGK promoter for expression of the tCD19 tag because the PGK promoter has been shown to not exhibit strong enhancer effects in studies comparing the activity of different promoters^33^. The CFTR cDNA was codon diverged to remove the sequence corresponding to the sgRNA in exon 1 and reduce homology to the endogenous sequence. The CFTR cDNA was split at 2883 bp (Amino acid 961, Methionine) from the ATG site. The sgRNA sequence from exon 1 was engineered into the first half of the HR template and this was followed by a 400 bp random stuffer sequence that does not bear homology to any human DNA sequence (Figure 2A). This stuffer serves as the right homology arm for the second HR template containing the remaining CFTR cDNA, a BGH polyA tail, a PGK promoter, truncated CD19 and an SV40 polyA tail.

**Fig. 2.**
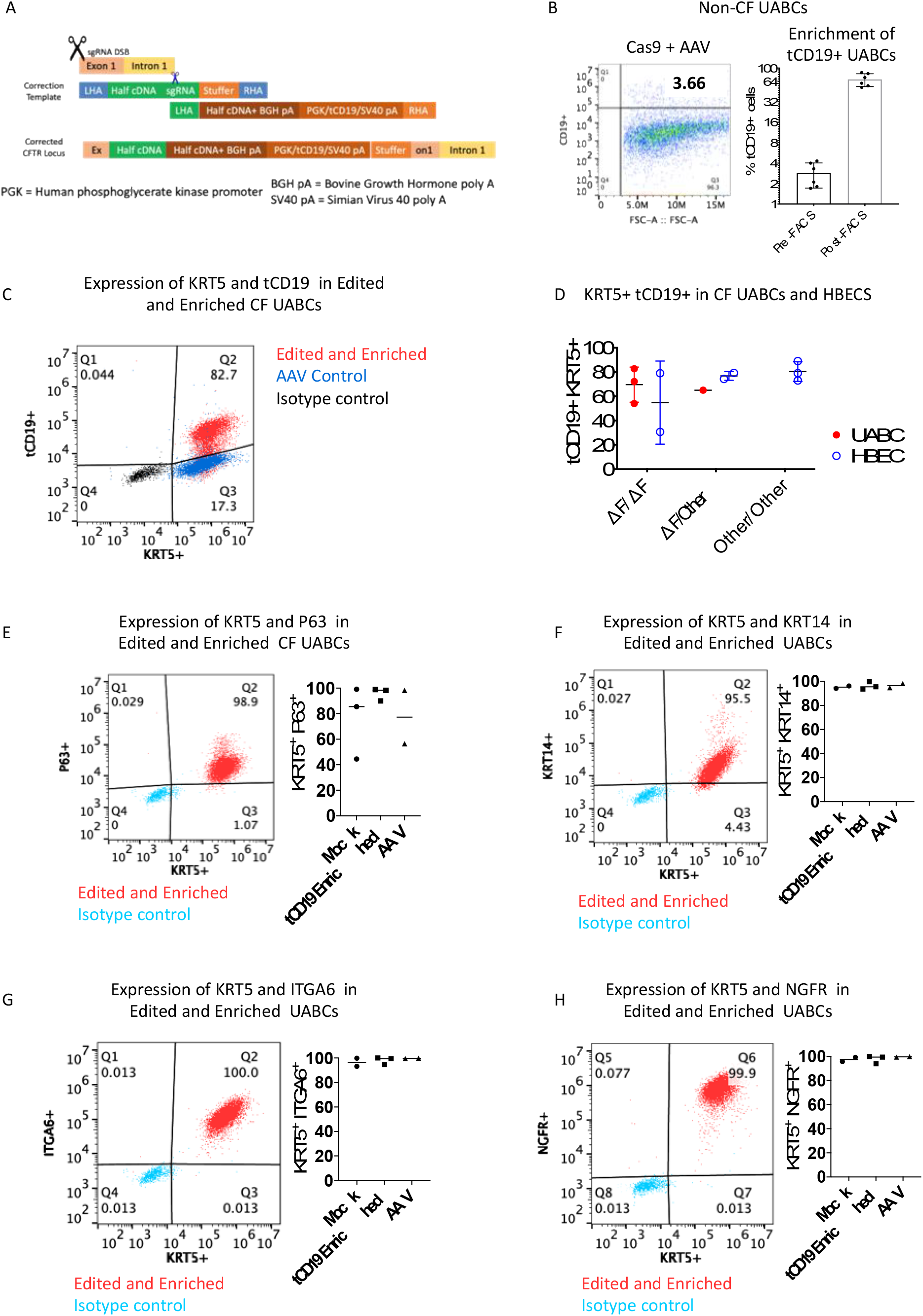
Insertion of the CFTR cDNA in UABCs and HBECs. (**A**) Schematic of the Universal strategy with the two halves of the CFTR cDNA and the tCD19 enrichment cassette. (**B**) After editing, 2-4% of edited UABCs were tCD19^+^. Edited UABCs were enriched by FACS to obtain >60% tCD19^+^ cells. A log base 2 scale has been used to make it easier to visualize the enrichment. (**C**) UABCs and HBECs from CF donors were edited using the Universal strategy. In this representative sample, 83% of the edited UABCs were positive for both tCD19 and KRT5. Controls treated with AAV but without Cas9 were KRT5^+^ and were negative for tCD19. **(D)** UABCs and HBECs from donors with different genotypes were corrected using the Universal strategy. In most cases >60% of enriched UABCs were tCD19^+^KRT5^+^. **(E)** >90% of FACS enriched UABCs were also positive for p63, **(F)** KRT14, **(G)** ITGA6 and **(H)** NGFR

Primary UABCs were electroporated with the Cas9/sgRNA complex and immediately incubated with two AAVs carrying the two halves of the HR template. We first tested this strategy in UABCs derived from donors without CF. We observed 3 ± 1% tCD19^+^ UABCs (Figure 2B). The edited UABCs were expanded for 10-14 days and tCD19^+^ cells were enriched using fluorescence activated cell sorting (FACS). We obtained 69 ± 19% tCD19^+^ cells after expansion of enriched cells (Figure 2B).

We extracted genomic DNA from the enriched tCD19^+^ cells to validate the proper insertion of the CFTR cDNA in the CFTR locus. The CFTR cDNA was codon-diverged from the native sequence in order to prevent premature recombination crossovers and to distinguish it from the endogenous sequence. We analyzed the presence of this codon-diverged CFTR sequence by Sanger-sequencing. The CFTR locus was amplified using junction-spanning primers at the 5′ insertion site, 3′ insertion site and at the junction in which the two halves of the CFTR cDNA are joined. We confirmed that the two halves of the CFTR cDNA were joined correctly by sequential HR (Figure S1B-C). We also amplified the beginning and the end of the CFTR cDNA insert and the tCD19 enrichment cassette and confirmed the presence of the expected sequences.

To determine the number of alleles in the pool of cells with our CFTR cDNA insert, we optimized a droplet digital PCR (ddPCR) assay. Figure S1D plots the percentage of tCD19^+^ cells observed with respect to the percent of alleles positive for the tCD19 sequence. The percentage of alleles modified as measured by ddPCR is approximately half the percentage of tCD19^+^ cells detected by FACS. This ratio is consistent with the insertion of the full length CFTR cDNA and tCD19 cassette into one allele per tCD19^+^ cell.

### Insertion of the CFTR cDNA into UABCs and HBECs obtained from donors with CF

After optimizing the gene correction and enrichment strategy on healthy donor UABCs, we tested the sequential insertion strategy in UABCs and HBECs derived from 11 different CF patients. UABCs were derived from 4 different CF patients who were all homozygous or compound heterozygous for F508del. In these 4 samples, we observed 4 ± 2% tCD19^+^ UABCs (Figure S1E) after editing. We started by gene editing 500,000 to 1 million UABCs (passage 1) and obtained 10 – 46 million UABCs within two passages post-editing (passage 3) (Figure S1F). Only 3 – 5 million of the expanded UABCs were used for further FACS enrichment. On average, about 100,000 ± 50,000 tCD19^+^ cells were obtained after FACS. These were expanded to obtain 500,000 – 2 million cells after expansion for 1 passage. If all the corrected cells had been FACS sorted, we estimate that up to 38 million cells could have been obtained after enrichment and expansion for one passage (Figure S1F). We obtained 68 ± 12% tCD19^+^ UABCs that were also KRT5^+^ after FACS enrichment and expansion in culture for 4 days (Figure 2C-D). On day 4 after FACS enrichment, >90% of the corrected UABCs were also positive for KRT5, P63, cytokeratin 14 (KRT14), integrin-alpha 6 (ITGA6) and nerve growth factor receptor (NGFR) (Figure 2E-H).

In order to test the ability of our strategy to restore CFTR function in patients with other genotypes including Class I nonsense mutations such as W1282X and R553X, we obtained HBECs derived from 7 different donors from 4 different sources. In HBECs derived from these 7 different donors with CF, we observed 10 ± 5% tCD19^+^ cells (Figure S1E) after editing and 72 ± 19% tCD19+ KRT5^+^ HBECs after enrichment (Figure 2D). Edited UABCs and HBECs stably express tCD19 only if both halves of the HR template are integrated into the genome. Episomal expression of tCD19 driven by the PGK promoter was measured using controls treated with only AAV and significantly fewer cells (p < 0.05) in the control group expressed tCD19 compared to edited samples at the time of enrichment (Figure S1E). Thus, we successfully inserted the cDNA in UABCs and HBECs from 11 different donors with CF and enriched using the tCD19^+^ cell surface expression to identify the cells with the full CF cDNA inserted.

### Restoration of CFTR function in CF Patient Samples

The restoration of CFTR function was tested on epithelial sheets generated by culturing the edited and enriched UABCs and HBECs at air-liquid interface (ALI) ^23^. UABCs differentiated in ALI cultures for 28-35 days produced differentiated epithelial sheets containing ciliated cells expressing acetylated alpha-tubulin and secretory cells expressing MUC5AC oriented toward the apical surface (Figure 3A). Moreover, the transepithelial resistances of differentiated epithelial sheets produced by UABCs cultured on BME and iMatrix maintained high transepithelial resistances, indicative of the production of a well-differentiated epithelium (Figure 3B). The percent of tCD19+ alleles in the edited and enriched cells were not significantly different before and after differentiation (paired t-test, p = 0.29) indicating that the corrected UABCs did not have a proliferative disadvantage (Figure 3C).

**Fig. 3.**
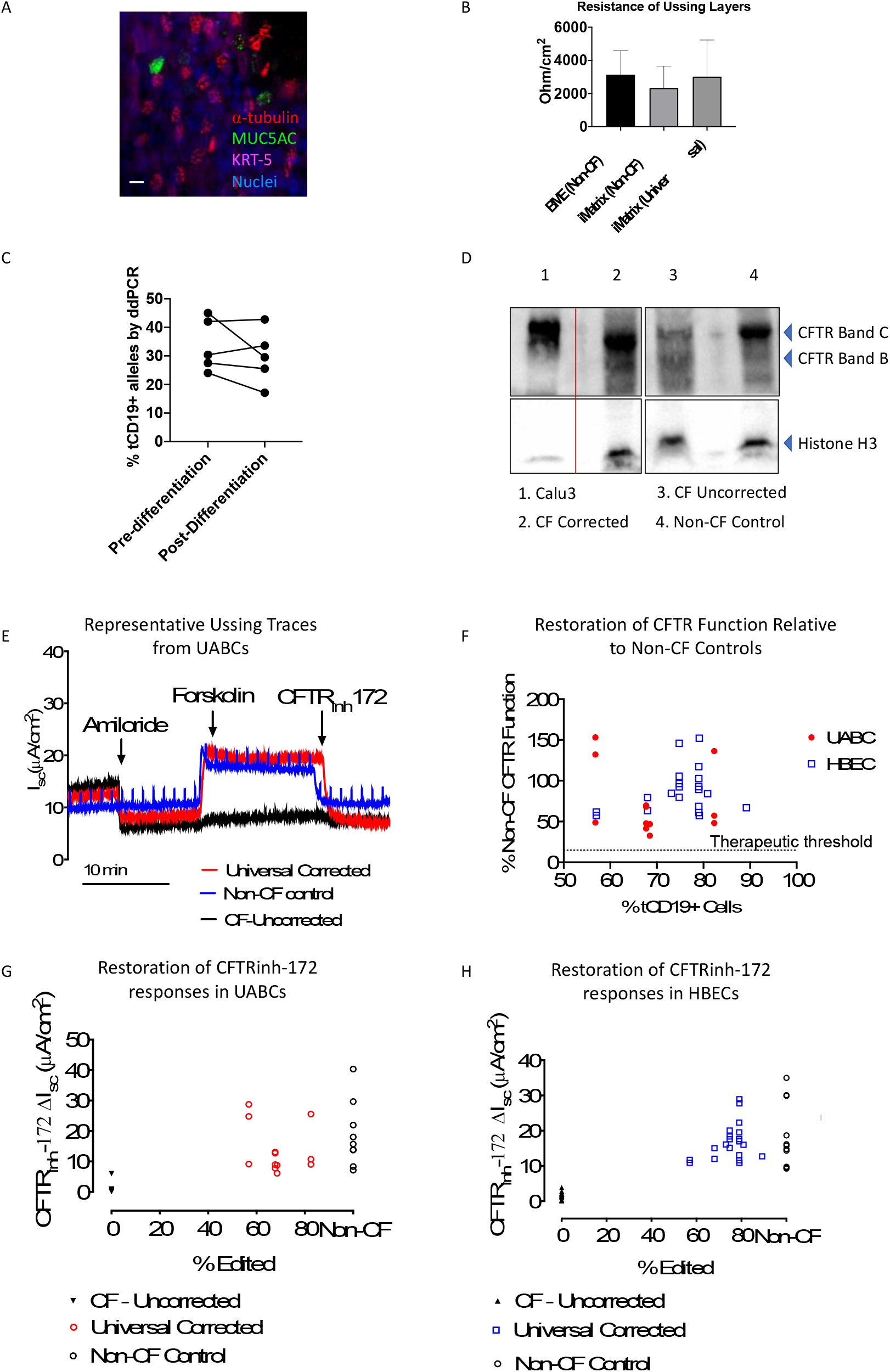
Restoration of CFTR expression and function in differentiated epithelial sheets. (**A**) Epithelial sheets generated by differentiation of UABCs (cultured on iMatrix) in air-liquid interfaces (ALI) contained cells positive for KRT5, acetylated tubulin and Mucin 5AC. (**B**) Epithelial sheets generated from UABCs cultured on iMatrix (unedited and Universal edited) showed trans-epithelial resistances that were not significantly different from epithelial sheets generated from UABCs cultured on BME. (**C**) The percent alleles modified was quantified both right after sorting and after differentiation for 28-35 days on ALI cultures. The percent of tCD19+ alleles was not significantly different between these timepoints indicating that the modified cells did not have any disadvantage in proliferation when compared to the unmodified tCD19-cells. (**D**) Western blot probing CFTR expression. CFTR expression in Calu-3 cells was used as a positive control (lane 1). Samples were diluted by 1/10 relative to other samples since Calu-3 cells are known to express high levels of CFTR. Lane 2 shows strong mature CFTR expression (band C) in a corrected CF sample, which is comparable to the mature CFTR expression (band C) observed in a non-CF control shown in lane 4. Uncorrected cells (F508del/3659delC) obtained from the same donor as the corrected cells used in lane 2 show reduced mature CFTR expression (band C, lane 3). (**E**) Representative traces obtained from epithelial sheets by Ussing chamber analysis in one non-CF control sample and one CF sample from donor 1 before and after correction. The traces from the corrected sample show similar forskolin and CFTR_inh-_172 responses as the non-CF control sample. (**F**) CFTR_inh-_172 short-circuit currents observed in corrected CF samples from 11 different donors as a percentage of the average CFTR_inh-_172 currents observed in non-CF controls. (**G**) CFTR_inh-_172 short-circuit currents observed in non-CF controls and epithelial monolayers derived from CF UABCs before and after correction. (**H**) CFTR_inh-_172 short-circuit currents observed in non-CF controls and epithelial monolayers derived from CF HBECs before and after correction.

We first measured CFTR expression in ALI cultures using immunoblotting. Figure 3D shows a western blot probing CFTR expression using anti-CFTR primary antibody Ab450 in non-CF, uncorrected, and corrected CF samples. The non-CF control shows one band corresponding to fully glycosylated mature CFTR (band C, 170-180 kDa) and incompletely glycosylated CFTR (band B) (Lane 4). The CF sample shows a weak band C (Lane 3) indicating impaired glycosylation of CFTR. The sample used in this study is a compound heterozygous sample (F508del/3659delC) with a frameshift present close to the end of the CFTR protein. The corrected sample showed an increase in band C intensity, demonstrating an increase in mature CFTR protein, relative to the uncorrected CF sample (Lane 2).

CFTR function was quantified by measuring short-circuit currents from ALI differentiated epithelial sheets. Representative traces obtained from an Ussing chamber assay for a CF sample before and after correction are shown in Figure 3E. Consistent with the observation of restored mature CFTR expression seen in the western blot, we observed restoration of CFTR function as indicated by the restoration of both the forskolin-stimulated short-circuit current and the inhibitable short-circuit current responsive to the selective CFTR inhibitor CFTR_inh_-172. The magnitudes of the responses to forskolin and CFTR_inh_-172 are similar to those observed in epithelial sheets derived from non-CF control donors. The CFTR_inh_-172 responses observed in corrected epithelial sheets derived from 11 different CF patients are presented in Table 1.

**Table 1:** Summary of Percent Editing of CF UABCs and HBEC Samples and Relative Restoration of CFTR Function as measured by responses to CFTR_inh_-172. CFTR contains 1480 amino acids. The mutations presented here occur in the first 1/3 or the last 1/3 of the CFTR protein. △F refers to the F508del mutation.

| Patient | Genotype | Editing Efficiency (% tCD19+ cells) | Raw Inhibitable CF Current $\mu\text{A}/\text{cm}^2$ | Percentage of Non-CF Inhibitable Current |
| --- | --- | --- | --- | --- |
| **Upper Airway Basal Cells (UABC)** |  |  |  |  |
| **WT average: $-19 \pm 11 \mu\text{A}/\text{cm}^2$** |  |  |  |  |
| **1** | $\Delta\text{F}/\Delta\text{F}$ | 67.7 | -13.1 | 69.81 |
| **1** | $\Delta\text{F}/\Delta\text{F}$ | 67.7 | -12.78 | 68.10 |
| **1** | $\Delta\text{F}/\Delta\text{F}$ | 67.7 | -8.96 | 47.75 |
| **1** | $\Delta\text{F}/\Delta\text{F}$ | 67.7 | -7.8 | 41.57 |
| **2** | $\Delta\text{F}/3659\text{delC}$ | 56.8 | -9.15 | 48.76 |
| **2** | $\Delta\text{F}/3659\text{delC}$ | 56.8 | -24.8 | 132.16 |
| **2** | $\Delta\text{F}/3659\text{delC}$ | 56.8 | -28.74 | 153.15 |
| **3** | $\Delta\text{F}/\Delta\text{F}$ | 68.5 | -8.79 | 46.84 |
| **3** | $\Delta\text{F}/\Delta\text{F}$ | 68.5 | -6.14 | 32.72 |
| **4** | $\Delta\text{F}/\Delta\text{F}$ | 82.3 | -25.59 | 136.37 |
| **4** | $\Delta\text{F}/\Delta\text{F}$ | 82.3 | -10.75 | 57.29 |
| **4** | $\Delta\text{F}/\Delta\text{F}$ | 82.3 | -9.04 | 48.17 |

| Human Bronchial Epithelial Cells (HBEC) |  |  |  |  |
| --- | --- | --- | --- | --- |
| WT average: $-18 \pm 9 \mu\text{A}/\text{cm}^2$ | | | | |
| 5 | $\Delta F/\Delta F$ | 57.0 | -10.9 | 57.31 |
| 5 | $\Delta F/\Delta F$ | 57.0 | -11.72 | 61.62 |
| 6 | $\Delta F/\Delta F$ | 79.0 | -22.31 | 117.29 |
| 6 | $\Delta F/\Delta F$ | 79.0 | -16.99 | 89.32 |
| 6 | $\Delta F/\Delta F$ | 79.0 | -19.53 | 102.68 |
| 6 | $\Delta F/\Delta F$ | 79.0 | -17.63 | 92.69 |
| 7 | $\Delta F/\text{N1303K}$ | 74.8 | -17.6 | 92.53 |
| 7 | $\Delta F/\text{N1303K}$ | 74.8 | -18.36 | 96.53 |
| 7 | $\Delta F/\text{N1303K}$ | 74.8 | -20.07 | 105.52 |
| 7 | $\Delta F/\text{N1303K}$ | 74.8 | -15.14 | 79.60 |
| 7 | $\Delta F/\text{N1303K}$ | 74.8 | -27.75 | 145.89 |
| 8 | 1717-1G/N1303K | 79.1 | -28.94 | 152.15 |
| 8 | 1717-1G/N1303K | 79.1 | -10.9 | 57.31 |
| 9 | $\Delta F/\text{W1282X}$ | 79.0 | -11.61 | 61.04 |
| 9 | $\Delta F/\text{W1282X}$ | 68.0 | -12.04 | 63.30 |
| 9 | $\Delta F/\text{W1282X}$ | 79.0 | -13.06 | 68.66 |
| 9 | $\Delta F/\text{W1282X}$ | 68.0 | -15.07 | 79.23 |
| 10 | W1282X/W1282X | 89.2 | -12.75 | 67.03 |
| 10 | W1282X/W1282X | 80.9 | -16.02 | 84.2239664 |
| 11 | W1282X/R553X | 73.1 | -16.1 | 84.6445604 |

Epithelial sheets derived from corrected UABCs and HBECs obtained from the 11 different donors with CF showed CFTR_inh_-172 responses that were 30-150 % of the CFTR_inh_-172 responses seen in non-CF controls (Table 1, Figure 3F). Corrected CF UABC cultures showed CFTR_inh_-172-sensitive short-circuit currents of 14 ± 8 μA/cm^2^ (74 ± 42% relative to non-CF) which was significantly higher than currents of 1 ± 2 μA/cm^2^ seen in uncorrected CF controls (adjusted p-value = 0.0085, ANOVA followed by Tukey test). Non-CF UABC cultures showed CFTR_inh_-172-sensitive short-circuit currents of 19 ± 11 μA/cm^2^ which is not significantly different from the CFTR_inh_-172 responses seen in corrected UABC cultures (adjusted p-value = 0.3540) (Figures 3F-G). Corrected CF HBEC cultures showed CFTR_inh_-172-sensitive short-circuit currents of 17 ± 5 μA/cm^2^ (88 ± 27% relative to non-CF) which was significantly higher than currents of 2 ± 1 μA/cm^2^ seen in uncorrected CF controls (adjusted p-value < 0.0001, ANOVA followed by Tukey test). This indicates the ability of the Universal strategy to restore CFTR function in samples with different genotypes (Table 1). Non-CF HBEC cultures showed CFTR_inh_-172-sensitive short-circuit currents of 18 ± 9 μA/cm^2^ which is not significantly different from the CFTR_inh_-172 responses seen in corrected UABC cultures (adjusted p-value = 0.6989) (Figures 3F, 3H).

### Off-target activity of sgRNA

The activity of Cas9 is directed to a target location in the genome by means of the sgRNA. However, Cas9 can tolerate mismatches between the sgRNA and the genomic DNA resulting in potential off-target activity. We used in-silico methods to identify over 50 possible off-target (OT) sites and characterized the activity of the sgRNA at the top 47 predicted off-target sites (Table S1) using both wild-type Cas9 (WT-Cas9) and high-fidelity Cas9 (HiFi Cas9), which was previously reported to be effective in maintaining high on-target activity while lowering off-target activity ^34^ (Figure S3, Table S1). We observed significant off-target activity in OT-3. We observed INDELs in 57 ± 3% alleles when we used WT-Cas9. However, this off-target activity was reduced by >30-fold to 1.2 ± 0.4% INDELs by using HiFi Cas9. Consistent with previous reports, we did not observe any reduction in on-target activity as measured by INDELs (Figure S3A)^34^. The off-target site (OT-3) corresponds to an intergenic region that has no known function and is located 10 kbp away from the nearest gene which codes for sideroflexin 1 which is a mitochondrial serine transporter. Table S1 presents the sequences and genes associated with each of the 47 off-target sites.

### No enrichment of edited cells with mutations in genes associated with solid tumors

Studies have reported that the ex vivo editing of cells may result in selection against cells with a functional p53 pathway and thus increase the risk of tumor formation if these cells were used for therapy ^35,36^. To characterize the enrichment of cells with increased tumor forming potential, we used the Stanford Solid Tumor Actionable Mutation Panel (STAMP) to detect the enrichment of clinically actionable mutations in 130 genes associated with the formation of solid tumors in edited and enriched UABCs obtained from 1 donor with CF and HBECs obtained from 2 different donors ^37–40^. No sample showed any mutation in *TP53* with or without editing. Mutations were only observed in 1-3 genes out of the 130 genes listed in the panel (Table S2). However, these mutations were also present in the control unedited samples from the same donors and the variable allele frequency (VAF) was not different between the control and edited samples. Because the identical mutations were present in both the edited and unedited samples, it means that the mutations must have been present in the cells originally (the shared common ancestor) and demonstrates that the engineering process did not generate them.

## Discussion

Several previous studies have attempted to add a functional copy of the CFTR cDNA using viral and non-viral methods both in vitro and in vivo ^6–8,13,14,41,42^. In these reports, the expression of CFTR was driven by exogenous promoters and the CFTR expression cassette was not inserted in the endogenous CFTR locus. Nevertheless, in vitro studies in human samples and an in vivo study in a CF pig model observed restoration of CFTR function following gene replacement ^13,14^. The development of CRISPR/Cas9 mediated genome editing enables the insertion of the CFTR cDNA in the endogenous locus and thus should enable the preservation of native CFTR expression, including in ciliated cells and ionocytes. However, it was unknown if the addition of the full-length CFTR cDNA in the endogenous locus under the control of the native promoter would be sufficient to restore CFTR function. Since the CFTR cDNA with the appropriate homology arms cannot be packaged into one AAV capsid, insertion of partial CFTR cDNAs into introns 7 and 8 of the CFTR gene has been attempted and this approach that can potentially be used to treat ∼89% of CF patients ^43^. This approach which did not enrich for corrected cells was tested in four CF patient samples and resulted in CFTR function that was 30-50% relative to non-CF controls. Our results show that the insertion of the full length CFTR cDNA which can treat >99.5% of CF patients results in CFTR function similar to that seen in non-CF controls (Figure 3F-H).

Concerns about the restoration of CFTR function using this Universal strategy stem from previous studies that reported the importance of introns in regulating CFTR expression ^44,45^. Our results show that insertion of the CFTR cDNA in exon 1 of the endogenous locus can successfully restore CFTR function in airway epithelial sheets generated from the UABCs and HBECs of CF patients. The successful restoration of CFTR might be enabled by the fact that all of the exons and introns downstream of exon 1 are left intact when cells are corrected using the Universal strategy. This result is also consistent with another study which showed that the insertion of a partial CFTR cDNA construct into intron 8 did not disrupt the open chromatin profile of the CFTR locus ^43^. Future studies characterizing the open chromatin profile of corrected UABCs and HBECs will further inform us about the influence of our Universal strategy on the regulation of both CFTR and neighboring genes.

Although our therapeutic approach is applicable to all CF patients harboring mutations in any combination of exons or introns, it is particularly relevant for patients with mutations that do not respond to recently developed CFTR modulators. These patients often have Class I nonsense mutations such as W1282X and exhibit a very severe disease burden and would not be responsive to current CFTR modulator therapies. Significantly, our experiments demonstrate the feasibility of restoring CFTR function in the presence of undruggable nonsense mutations such as W1282X and R553X (Table 1). Our approach thus enables the development of an autologous airway stem cell therapy for all CF patients affected by mutations in any of the exons or introns of the CFTR gene including patients with nonsense mutations. Additional strategies may be needed to treat patients with biallelic mutations in the promoter, estimated to be <0.5% of the CF population.

Our experiments further show that the corrected UABCs and HBECs produce differentiated epithelial sheets with restored CFTR function (Figure 3). In epithelial sheets produced from corrected UABCs and HBECs obtained from 11 different CF patients, we measured restored CFTR mediated chloride transport that was on average >70% of the levels seen in non-CF controls with individual readings ranging from 32% - 150% of control values. Several previous studies have shown that when CF cells were either mixed with cells transduced with viruses inducing the over-expression of CFTR ^42,46–48^ or wild type cells ^49,50^, the presence of 10-25% CFTR-expressing cells was sufficient to restore CFTR function close to levels in non-CF controls. Other studies have investigated CFTR function in patients with milder CF phenotypes associated with improved survival and these studies also observed as little as 2-15% CFTR function relative to wild-type ^51,52^. Our studies indicate that the Universal strategy can restore CFTR function well beyond this 15% threshold and enables the restoration of CFTR function to levels seen in asymptomatic CF carriers ^51,52^.

In addition to validating the Universal CFTR correction strategy, our studies also advance our ability to generate a clinical cell therapy product by replacing xenobiotic reagents such as BME with reagents compatible with good manufacturing practices. We previously reported the feeder-free culture of UABCs in a chemically defined media formulation ^23^. However, the UABCs were plated on tissue culture treated plates coated with BME derived from mouse sarcoma cells. Previous studies have reported the improved expansion of airway basal stem cells using laminin enriched conditioned media along with dual-SMAD inhibition ^53^. Improving upon these reports, the results from this current study indicate that recombinant laminin 511 (iMatrix™) is an effective replacement for BME that further improves our ability to expand UABCs using reagents compatible with the production of airway stem cells for clinical applications. Moreover, the primary UABCs cultured in this manner do not experience a significant shortening of their telomeres, presumably due to the retention of some telomerase activity (Figure 1 C-E). Telomerase activity has been reported to be associated with the self-renewal potential of hepatocytes and hematopoietic stem cells ^54,55^. In addition, telomerase activity has also been reported in airway cells obtained from freshly isolated nasal tissue ^56^. Thus, our results suggest that the durability of the primary UABCs used in our studies is unlikely to be limited by shortened telomeres. Lastly, previous studies in hematopoietic cells have reported that efficient HR is facilitated by improved cell proliferation ^27^ and these results were also consistent with our previously reported findings that improved culture conditions facilitate HR in airway stem cells ^23^. In this study, we did not evaluate if the modified culture conditions improved HR significantly.

In addition to assessing the potential durability of the gene corrected stem cells, it is also important to test their safety. Previous studies that reported the enrichment of cells with dysfunctional *TP53* pathway used immortalized cells or primary cells from model organisms such as mice ^35,36^. Here, we have quantified the presence of mutations in *TP53* and an additional 129 genes associated with solid tumors in gene corrected primary UABCs and HBECs from humans. We did not observe any increase in the frequency of alleles with mutations in genes associated with solid tumors. These results are consistent with previous reports from our group which did not observe any enrichment of alleles with mutations in *TP53* in gene edited hematopoietic stem and progenitor cells and iPSCs ^37,57^. Thus, our results indicate that our Universal ex vivo strategy for CFTR may be safe for further clinical translation.

We previously reported our ability to embed corrected UABCs on a porcine small intestinal submucosal (pSIS) membrane that is FDA approved for use in facilitating sino-nasal repair after surgeries ^23^. In that previous study, we reported the need for 1.3 million UABCs in order to fully cover one maxillary sinus ^23^. The use of the optimized culture protocols reported here enable us to obtain this yield (Figure S1F). These studies thus build the foundation for the development of an autologous gene corrected stem cell therapy for all CF patients. The transplantation of gene corrected airway stem cells and the restoration of mucociliary transport in vivo is still a significant hurdle that must be overcome for the clinical translation of this approach. Future studies will first investigate the transplantation of UABCs embedded on the pSIS membrane or other biomaterials into the sinuses of animal models to treat CF sinus disease, an important unmet need and common morbidity in nearly all CF patients. The knowledge developed in this process will be useful in optimizing the autologous transplantation of gene corrected airway basal stem cells into the lower airways as a durable stem cell therapy for CF lung disease.

## Conclusion

Our experiments show that the full-length CFTR cDNA along with an enrichment cassette expressing tCD19 can be inserted into the endogenous CFTR locus in airway basal stem cells using Cas9 and two AAV6 vectors containing two halves of the HR template. The corrected airway basal stem cells differentiate to produce epithelial sheets that exhibit restored CFTR mediated chloride transport that is on average 70% - 80% of the levels seen in non-CF controls. Our findings enable the further development of a durable therapeutic option for all CF patients, especially those patients who cannot be treated using existing modulator therapies. In addition, the findings enable future studies to optimize the transplantation of gene corrected UABCs into the upper airway epithelia of CF patients to treat CF sinus disease, as well as strategies to develop an autologous gene corrected airway stem cell therapy for CF lung disease.

## Materials and Methods

### Subject details

Upper airway tissue was obtained from adult patients undergoing functional endoscopic surgery (FESS) after obtaining informed consent. The protocol was approved by the Institutional Review Board at Stanford University. The CFTR mutations from each subject were recorded. HBECs were obtained from Lonza Inc., the Primary Airway Cell Biobank at McGill University, the Cystic Fibrosis Foundation Cell Bank or the University of North Carolina. CF bronchial epithelial cells were obtained from lungs explanted during transplantation under protocol #13-1396 approved by the Committee on the Protection of the Rights of Human Subjects at the University of North Carolina at Chapel Hill. All donors provided informed consent.

## METHOD DETAILS

### EN media

ADMEM/F12 was supplemented with B27 supplement, Nicotinamide (10 mM), human EGF (50 ng/mL), human Noggin (100 ng/mL), A83-01 (500 nM), N-acetylcysteine (1 mM) and HEPES (1 mM)

### Cell Culture of UABCs

Upper airway tissue pieces were cut into small pieces (1-2 mm^2^). Tissue pieces were washed with 10 mL sterile PBS with 2X antibiotic/antimycotic (Penicillin, Streptomycin, Amphotericin B, Gibco #15240062) on ice and digested with pronase (1.5 mg/mL, Sigma #P5147) for 2 hours at 37 °C or at 4°C overnight. Digestion was stopped using 10% fetal bovine serum (FBS). Digested tissue was filtered through cell strainers (BD Falcon #352350) into a sterile 50 mL conical tube. The mixture was centrifuged at 600 x g for 3 minutes at room temperature. Red blood cell (RBC) lysis was then performed using RBC lysis buffer (eBioscience™) as per manufacturer’s instructions. After RBC lysis, cells were suspended in 1 mL EN media and counted. A small sample was fixed using 2% paraformaldehyde and permeabilized using Tris-buffered saline with 0.1% Tween 20. Cells were stained for cytokeratin 5 (KRT5, Abcam, ab 193895). An isotype control (Abcam, ab 199093) was used to assess non-specific staining. KRT5^+^ cells were plated at a density of 10,000 cells per cm^2^ as monolayers in tissue culture plates. At the time of plating, 4.5 μL of iMatrix™ per well of a 6 well plate was added immediately after the addition of EN media into the wells. Cells were incubated at 37°C in 5% O_2_and 5% CO_2_in EN media with 10 μM ROCK inhibitor (Y-27632, Santa Cruz, sc-281642A). Cells obtained from CF patients were grown in EN media supplemented with additional antimicrobials for two days (Fluconazole – 2 μg/mL, Amphotericin B 1.25 μg/mL, Imipenem – 12.5 μg/mL, Ciprofloxacin – 40 μg/mL, Azithromycin – 50 μg/mL, Meropenem - 50 μg/mL). The concentration of antimicrobials was decreased by 50% after 2-3 days and then withdrawn after editing (day 5-6).

### Cell Culture of HBECs

Human bronchial epithelial cells were cultured in Pneumacult™ Ex-Plus at 3,000-10,000 cells/cm^2^ in tissue culture flasks without any coating.

### Gene Editing of UABCs

UABCs and HBECs were edited as previously reported but with sgRNA and HR templates specific to the Universal correction strategy ^23^. Briefly, cells were cultured in EN media with 10 μM ROCK inhibitor (Y-27632, Santa Cruz, sc-281642A). Gene correction was performed 5 days after plating. Cells were detached using TrypLE Express Enzyme (Gibco™ 12605010). 5 × 10^5^ – 1 × 10^6^ cells were resuspended in 100 μL OPTI-MEM (Gibco™ 31985062) resulting in a density of 5-10 million cells/mL. Electroporation (Nucleofection) was performed using Lonza 4D Nucleocuvette™ vessel (Lonza, V4XP-3024). 30 μg of WT-Cas9 (Integrated DNA Technologies, IA, Cat: 1074182) or HiFi Cas9 (Integrated DNA Technologies, IA, Cat:1081061) and 16 μg of MS-sgRNA (Trilink Biotechnologies, CA) (molar ratio = 1:2.5) were complexed at room temperature for 10 minutes, mixed with the cells suspended in 100 μL of OPTI-MEM and transferred to a Nucleocuvette™ vessel. Cells were electroporated using the program CA-137. 400 μL of OPTI-MEM was added to each well after electroporation. Two AAVs carrying the two halves of the CFTR cDNA and the tCD19 enrichment cassette were added at a Multiplicity of Infection (MOI) of 10^5^ genomes per cell (as determined by ddPCR). Our previous study reported an optimal MOI of 10^6^ genomes per cell. However, those experiments used qPCR to quantify AAV genomes which overestimates the number of AAV genomes. Functional titration experiments were used to determine the optimal AAV MOIs for this study. Cells along with the AAVs were transferred into multiple wells of a 6-well plate such that the cell density was ∼5,000 cells/cm^2^. 1.5 mL of EN media followed by 4.5 μL of iMatrix™ were added to each well. Media was replaced 48 hours after electroporation.

UABCs were passaged 4-5 days after editing in order to expand the edited cells for FACS enrichment. Passaged UABCs were plated in tissue culture treated flasks with a surface area of 225 cm^2^. 100 μL of iMatrix™ was added to each flask in addition to 25 mL of EN media. UABCs were suspended by trypsinization 4-5 days after passaging and enriched for tCD19^+^ cells using FACS. tCD19 was detected using an anti-human CD19 antibody (Biolegend, 302206)

### Gene Editing of HBECs

The same protocol used for UABCs was used for HBECs except that the cells were cultured in Pneumacult™ Ex-Plus supplemented with 10 μM ROCK inhibitor.

### FACS enrichment of edited UABCs and HBECs

Edited UABCs and HBECs were suspended by treatment with TrypLE™ Express Enzyme (Gibco™ 12605010). UABCs were incubated with anti-human-CD19 antibodies conjugated with FITC for 20-30 minutes. Cells were also incubated with 7-AAD for 10 minutes in order to label non-viable cells. The labeled cells were washed three times with FACS buffer (PBS with 1% BSA and EDTA) and resuspended in FACS buffer. The UABCs and HBECs were FACS sorted using Aria II SORP (BD Biosciences). For each experiment, cells that were unmodified and cells that were treated with only the AAVs but no Cas9 were used as controls in order to identify FACS settings that specifically detected genome edited tCD19^+^ UABCs and HBECs.

### Measuring Gene Correction using analytical flow cytometry

Cells were expanded for 4-5 days after FACS sorting. The percent of tCD19^+^ cells were quantified using a FACS analyzer (Beckman Coulter, CytoFLEX). Cells were also stained for other basal stem cell markers such as cytokeratin 14 (KRT14, Abcam, ab181595), P63 (Biolegend, 687202), Integrin Alpha 6 (ITGA6, Biolegend, 313608) and Nerve Growth Factor Receptor (NGFR, Biolegend, 345106).

### Measuring Gene Correction using ddPCR

Correction was also validated using droplet digital PCR. Genomic DNA was extracted from cells using Quick Extract (QE, Lucigen, QE09050). The cells were suspended by trypsinization and washed once with OPTI-MEM or media. 50-100,000 cells were centrifuged, and the supernatant was discarded. 50μL quick extract (QE) was added to the cell pellet and vortexed. The quick extract suspension was heated at 65°C for 6 minutes, vortexed and then heated at 98°C for at least 10 minutes to inactivate QE. We observed that inactivation for less than 10 minutes often resulted in failed PCR. The modified CFTR locus was amplified using the primers:

Forward:AGCATCACAAATTTCACAAATAAAGCA

Reverse: ACCCCAAAATTTTTGTTGGCTGA

Probe sequence: CACTGCATTCTAGTTGTGGTTTGTCCA

A region in the intron 1 of CFTR was used as a reference for allele quantification using ddPCR. The reference region was amplified using:

Forward: TGCTATGCCAGTACAAACCCA

Reverse primer: GGAAACCATACTTCAGGAGCTG

Probe sequence: TTGTTTTGTATCTCCACCCTG

### Measuring off-target activity

Primary UABCs from non-CF patients were electroporated with Cas9 and sgRNA without the HR template. gDNA was extracted 4-5 days after RNP delivery. Potential off-target sites were identified using the bioinformatic tool COSMID ^58^ allowing for 3 mismatches within the 19 PAM proximal bases. Predicted off-target loci were initially enriched by locus specific PCR followed by a second round of PCR to introduce adaptor and index sequences for the Illumina MiSeq platform. All amplicons were normalized, pooled and quantified using a Qubit (ThermoFisher Scientific) and were sequenced using a MiSeq Illumina using 2 x 250bp paired end reads. INDELs at potential off-target sites were quantified as previously described ^59^.

Briefly, paired-end reads from MiSeq were filtered by an average Phred quality (Qscore) greater than 20. Single reads were quality trimmed using Cutadapt and Trim Galore prior to merging of paired end reads using Fast Length Adjustment of SHort reads (FLASH). Alignments to reference sequences were performed using Burrows-Wheeler Aligner for each barcode. Percentage of reads containing insertions or deletions with a ± 5-bp window of the predicted cut sites were quantified.

### Air-Liquid Interface Culture of Corrected UABCs and HBECs

Gene corrected cells were plated either on the day of FACS sorting or after expansion for 4-5 days. 30,000 to 60,000 cells per well were plated in 6.5 mm Transwell plates with 0.4 μm pore polyester membrane insert (Corning Inc., 07-200-154). EN media was used to expand cells until they were confluent (3-7 days). Once cells were confluent in Transwell inserts and stopped translocating fluid between the upper and lower chambers, media in the bottom compartment was replaced with UNC ALI media previously reported by Gentzsch et al ^60^.

### Immunoblot

Immunoblotting methods were used to compare CFTR protein expression pre/post-correction. Non-CF nasal cells, and patient-derived CF cells pre/post-correction were plated and cultured according to the above methods. Lysis was performed by incubating cells for 15 minutes in ice-cold RIPA buffer supplemented with EDTA-free protease inhibitor. Lysates were gathered and rotated at 4°C for 30 min, then spun at 10,000 × g at 4°C for 10 minutes to pellet insoluble genomic material. The supernatant was collected and mixed with 4x Laemmli sample buffer containing 100 mM DTT and subsequently heated at 37°C for 30 min. Lysate from ∼10,000 cells was loaded for Calu-3 lane, and lysates from ∼100,000 cells were loaded for non-CF nasal, uncorrected and corrected CF cells and fractionated by SDS-PAGE, then transferred onto PVDF membrane. Blocking was performed with 5% nonfat milk in TBST (10 mM Tris, pH 8.0, 150 mM NaCl, 0.5% Tween 20) for 60 min. The membrane was probed with antibodies against CFTR (Ab450, 1:1000), and Histone H3 (1:10,000). Membranes were washed and incubated with a 1:10,000 dilution of horseradish peroxidase-conjugated anti-mouse for 1 hour, then developed by SuperSignal™ West Femto Maximum Sensitivity Substrate (Thermo Fisher Scientific, 34095).

### Ussing Chamber Functional Assays

Ussing chamber measurements were performed 3-5 weeks after cells had stopped translocating fluid as described before ^23^. For chloride secretion experiments with UABCs and HBECs, solutions were as following in mM: Apical: NaGluconate 120, NaHCO_3_ 25, KH_2_PO_4_3.3, K_2_HPO_4_0.8, Ca(Gluconate)^2^ 4, Mg(Gluconate)^2^ 1.2, Mannitol 10; Basolateral: NaCl 120, NaHCO_3_, 25, KH_2_PO_4_ 3.3, K_2_HPO_4_ 0.8, CaCl_2_ 1.2, MgCl_2_ 1.2, Glucose 10The concentration of ion channel activators and inhibitors were as follows: Amiloride (10 μM, Apical), Forskolin (10 μM, Bilateral), VX-770 (10 μM, Apical), CFTR_inh_-172 (20 μM, Apical), UTP (100 μM, Apical).

### Protein, RNA, and genomic DNA extractions from single UABC pellets for telomerase and telomere measurements

Frozen cell pellets containing ∼ 1×10^6^ UABCs were resuspended in 100µl of NP40 lysis buffer (25mM HEPES, 150mM KCl, 1.5mM MgCl_2_, 0.5% NP40, 10% glycerol, PH adjusted to 7.5 with KOH, with freshly added 1 mM DTT, 200 µM PMSF, and 1:1000 Protease Inhibitor Cocktail [sigma, P8340]). Soluble supernatant was used for telomerase detection by TRAP (Telomeric Repeat Amplification Protocol), western blotting, as well as soluble RNA extraction; insoluble pellet is sufficient for genomic DNA (gDNA) extraction for telomere length measurement by Telo-qPCR and TRF (Telomere/Terminal Restriction Fragment) analyses.

For TRAP, 2µg of protein extract per Bradford were programed into a 25µl of “cold extension” reaction, 0.5µl from which was subsequently PCR-amplified in presence of p^32^-labeled TS primers, according to Chen et al.^61^ Total RNA was Trizol-extracted from the same soluble extract, and subjected to qPCR analysis for TERT mRNA expression, which tightly correlates with TRAP activity (data not shown).

For telomere length measurements, gDNA from the insoluble pellet was resuspended in tail lysis buffer containing Protease K, and then extracted by phenol-chloroform. 10ng of gDNA was used in Telo-qPCR reactions. Telomeric amplicon was PCR-amplified with 5′-ACACTAAGGTTTGGGTTTGGGTTTGGGTTTGGGTTAGTGT-3′, and 5′-TGTTAGGTATCCCTATCCCTATCCCTATCCCTATCCCTAACA-3′; Albumin housekeeping amplicon was amplified with 5′-CGGCGGCGGGCGGCGCGGGCTGGGCGGAAATGCTGCACAGAATCCTTG-3′, and 5′-GCCCGGCCCGCCGCGCCCGTCCCGCCGGAAAAGCATGGTCGCCTGTT-3′, according to Cawthon.^62^ For TRFs, 2µg of gDNA was digested with a restriction enzyme cocktail, containing *Hinf*I, *Rsa*I, *Msp*I, and *Alu*I, for 4 hours at 37°C. In-gel Southern hybridization was performed with a 3x TTAGGG p^32^-labelled probe, according to Herbert et al. ^63^

### Stanford Solid Tumor Actionable Mutation Panel

UABCs and HBECs obtained from donors with CF were corrected and enriched using FACS. The FACS enriched cells and control cells were then expanded for 1-2 passages to obtain ∼1 million cells per condition. The genomic DNA was extracted using GeneJET Genomic DNA Purification Kit (ThermoFisher Scientific, Cat: K0722).

Next generation sequencing of samples was performed at the Stanford Molecular Genetic Pathology Clinical Laboratory to quantify mutations in 130 genes (e.g. *TP53, EGFR, KRAS, NRAS*) identified in the Stanford Solid Tumor Actionable Mutation Panel (STAMP) ^38–40^. The STAMP assay is clinically validated and can detect variants with a variant allele fraction as low 5%.

The assay was performed as previously described to assess the enrichment of tumor forming mutations in gene edited hematopoietic cells using a similar panel ^37^. Briefly, the gDNA was sheared (M220 focused ultrasonicator, Covaris, Woburn, MA), sequencing libraries were prepared (KK8232 KAPA LTP Library Preparation Kit Illumina Platforms, KAPABiosystems, Wilmington, MA), and hybridization-based target enrichment was performed with custom-designed oligonucleotides (Roche NimbleGen, Madison, WI). Illumina sequencing instruments were used to sequence pooled libraries (MiSeq or NextSeq 500 Systems, Illumina, San Diego, CA). Sequencing results are analyzed with an in-house developed bioinformatics pipeline. Sequence alignment against the human reference genome hg19 is performed with BWA in paired end mode using the BWAMEM algorithm and standard parameters. Variant calling was performed separately for single nucleotide variants (SNVs), insertions and deletions < 20 bp (Indels), and fusions. VarScan v2.3.6 was used for calling SNVs and Indels, and FRACTERA v1.4.4 was used for calling fusions. Variants are annotated using Annovar and Ensembl reference transcripts.

## Acknowledgments

We appreciate the efforts of Ivan T. Lee, Nicole Borchard, Sachi Dholakia, David Zarabanda, Phillip Gall and Yasuyuki Noyama in prioritizing efforts at human tissue acquisition from the operating theater. We thank the CF Canada Primary Airway Cell Biobank at McGill University and the American Cystic Fibrosis Foundation Cell Bank for providing primary bronchial airway epithelial cells. CFTR antibodies were obtained from Dr. John Riordan at the University of North Carolina-Chapel Hill through the CF foundation antibody distribution program. Funding: The work was funded by grants from the California Institute of Regenerative Medicine (DISC2-09637), Stanford-SPARK MCHRI, NIH (K08DE027730 to A.A.S, U01DK085527, U01CA217851, U01CA176299 and U19AI116484 to C.J.K), the Cancer Prevention and Research Institute of Texas (RR14008 to G.B.) and the Cystic Fibrosis Foundation (BAO19XX0). We thank the Crandall Foundation for a philanthropic gift that supported this work. S.V and Z.M.S were supported by a fellowship from the Cystic Fibrosis Foundation (VAIDYA19F0, SELLER16L0). S.H.R acknowledges funding from the NIH (DK065988) and CF Foundation (BOUCHE19RO)

