## Supplemental Information for "Targeted replacement of full-length CFTR in human airway stem cells by CRISPR/Cas9 for pan-mutation correction in the endogenous locus"

**Supplementary Materials:**

Table S1: Predicted off target sites and the associated genes. Cas9 can induce double stranded breaks even if there are a few mismatches exist between the target genomic DNA and sgRNA. As a result, undesired insertions and deletions at off-target (OT) sites may occur. The potential off-target sites are presented along with the genes closest to the off-target sites.

| | Sequence | Description of the nearest genes |
|---|---|---|
| **On Target** | TTCCAGAGGCGACCTCTGCATGG | cystic fibrosis transmembrane conductance regulator |
| **OT1** | **C**TCCAG**GA**GCGACCTCTGCATGG | long intergenic non-protein coding RNA 293 |
| **OT2** | T**C**CC**T**G**G**GGCGACCTCTGCAGGG | long intergenic non-protein coding RNA 242 |
| **OT3** | TT**T**CAGAGGC**C**ACCTCTGCATGG | sideroflexin 1 |
| **OT4** | **G**TC**A**AGA**T**G**T**GACCTCTGCATGG | PR/SET domain 7 |
| **OT5** | **G**TC**A**AGA**T**G**T**GACCTCTGCATGG | PR/SET domain 9 |
| **OT6** | **G**T**G**CAGAGGCG**C**CCTCTGCAAAG | uncharacterized LOC100506585 |
| **OT7** | **G**T**TA**AGAGGC**T**ACCTCTGCAGGG | lysosomal protein transmembrane 5 |
| **OT8** | TTCC**GC**AGGC**A**ACCTCTGCATGG | sialic acid binding Ig like lectin 16 |
| **OT9** | **CA**CCAGA**C**GC**C**ACCTCTGCAAGG | uncharacterized LOC101929217 |
| **OT10** | **A**T**T**CAGA**T**GC**C**ACCTCTGCAGGG | kazrin, periplakin interacting protein |
| **OT11** | **CA**CCAGAG**C**C**C**ACCTCTGCAAGG | single-minded family bHLH transcription factor 2 |

| | | |
|---|---|---|
| **OT12** | TTCCAG**CA**GC**C**ACCTCTGCAAGG | NADPH oxidase 5 |
| **OT13** | **GC**CCAGAGGCGA**G**CTCTGCACAG | LIM homeobox 3 |
| **OT14** | **C**TCCAG**T**G**C**CG**T**CCTCTGCAGGG | smoothened, frizzled class receptor |
| **OT15** | **G**TCCAG**T**GG**TC**ACCTCTGCACGG | free fatty acid receptor 2 |
| **OT16** | TT**G**CAGAGG**A**G**C**CCTCTGCACGG | microRNA 4478 |
| **OT17** | **A**TC**T**AGAGG**A**G**G**CCTCTGCAGGG | zinc finger protein 664 |
| **OT18** | **C**TCCAGAGG**A**GA**T**CTCTGCAGAG | p21 protein Cdc42/Rac)-activated kinase 2 pseudogene |
| **OT19** | **A**TCCA**A**AGG**A**GA**G**CTCTGCAAGG | long intergenic non-protein coding RNA 1037 |
| **OT20** | **C**TCCAGAGGCG**GG**CTCTGCAGAG | uncoupling protein 3 |
| **OT21** | **AG**CCAGAGGC**C**A**G**CTCTGCAGGG | transglutaminase 4 |
| **OT22** | **C**T**G**CAGAGGC**C**AGCTCTGCATGG | uncharacterized LOC101927549 |
| **OT23** | **C**TCCAGAGG**G**GACC**C**CTGCAGAG | sushi domain containing 2 |
| **OT24** | T**C**CCAGAGGC**C**ACC**A**CTGCAGGG | ALG12, alpha-1,6-mannosyltransferase |
| **OT25** | **C**T**G**CAGAGGC**A**ACC**A**CTGCACGG | suppressor of cytokine signaling 1 |
| **OT26** | **A**TCCAGAGGC**TT**C**A**TCTGCATGG | POC1 centriolar protein A |
| **OT27** | **G**TCCAG**G**GGC**A**ACC**C**CTGCAGGG | uncharacterized LOC100507548 |
| **OT28** | **AA**CCAGAGGCG**T**CC**A**CTGCAGGG | long intergenic non-protein coding RNA 2133 |
| **OT29** | **C**TCCAGA**T**GCG**G**CC**C**CTGCAAGG | distal-less homeobox 4 |
| **OT30** | TTCCAGAGGC**A**A**GG**TCTGCAGGG | microRNA 548z |
| **OT31** | **G**TCCAGAGG**T**G**T**CC**C**CTGCAGGG | teashirt zinc finger homeobox 3 |
| **OT32** | **C**TCCAGAGGC**AG**CC**C**CTGCAGGG | glutamate metabotropic receptor 4 |
| **OT33** | TTCCAGAGG**GC**ACCT**T**TGCAAGG | anaphase promoting complex subunit 4 |

| | | |
|---|---|---|
| **OT34** | **C**TCCAGAGGC**TG**CCT**T**TGCAAGG | CCHC-type zinc finger nucleic acid binding protein |
| **OT35** | **C**TCCAGAGGC**C**AC**A**T**T**TGCAGGG | echinoderm microtubule associated protein like 4 |
| **OT36** | **C**TCCAGAGGC**TT**CCTC**A**GCAGGG | long intergenic non-protein coding RNA 523 |
| **OT37** | **C**TCCAG**G**GG**A**GACCTCT**T**CAGGG | uncharacterized LOC399715 |
| **OT38** | **G**TCCA**C**AGGCG**G**CCTCTTCATGG | kazrin, periplakin interacting protein |
| **OT39** | **G**TCCAGAGGC**CC**CCTCT**C**CAGGG | adenylate cyclase 2 |
| **OT40** | **G**TCCAGAGG**T**GACCTGT**C**CAAGG | immunoglobulin-like and fibronectin type III domain containing 1 |
| **OT41** | **C**TCCAGAGGC**T**AC**A**TCT**GG**AGGG | family with sequence similarity 9 member C |
| **OT42** | TTCCAGAGG**GA**ACCTCTGC**C**AGG | microRNA 149 |
| **OT43** | **A**TCC**G**GAGGCGACC**A**CTGC**C**TGG | G protein-coupled receptor 52 |
| **OT44** | TTCCAGAGG**T**GACCTCTTAATGG | dihydropyrimidine dehydrogenase |
| **OT45** | **C**TCCAGAGGCG**C**CCTCT**AG**AGGG | cholecystokinin B receptor |
| **OT46** | **A**TCCAGAGG**T**GACCTCT**C**C**C**AGG | proline dehydrogenase 1 |
| **OT47** | **G**TCCAGAG**C**CGACCTCT**GAG**GGG | C5orf66 antisense RNA 2 |

Table S2: Mutations in 130 genes associated with solid tumors were assessed using the STAMP panel. The table lists mutations observed with a variable allele frequency greater (VAF) than 5%. The patients were heterozygous for the listed mutations and the VAF was same between control and edited cells that were FACS enriched.

| | | | | | | | **Control** | **Edited and enriched** |
|---|---|---|---|---|---|---|---|---|
| **Cell type** | **Patient ID (Table 1)** | **Chr** | **Position** | **Gene** | **CDS Change** | **AA Change** | **VAF%** | **VAF%** |
| **UABC** | 2 | chr16 | 2112558 | TSC2 | c.1318G>A | p.G440S | 48 | 45.18 |
| **HBEC** | 7 | chr3 | 89259122 | EPHA3 | c.266G>A | p.R89K | 51.6 | 50.07 |
| | | chr1 | 16464805 | EPHA2 | c.944G>A | p.R315Q | 48.92 | 48.84 |
| | | chr7 | 55249005 | EGFR | c.2303G>A | p.S768N | 46.7 | 48.36 |
| **HBEC** | 9 | chr22 | 23634790 | BCR | 2845G>A | V949I | 50.67 | 49.47 |
| | | chr17 | 37883638 | ERBB2 | 3250G>T | D1084Y | 48.42 | 49.46 |

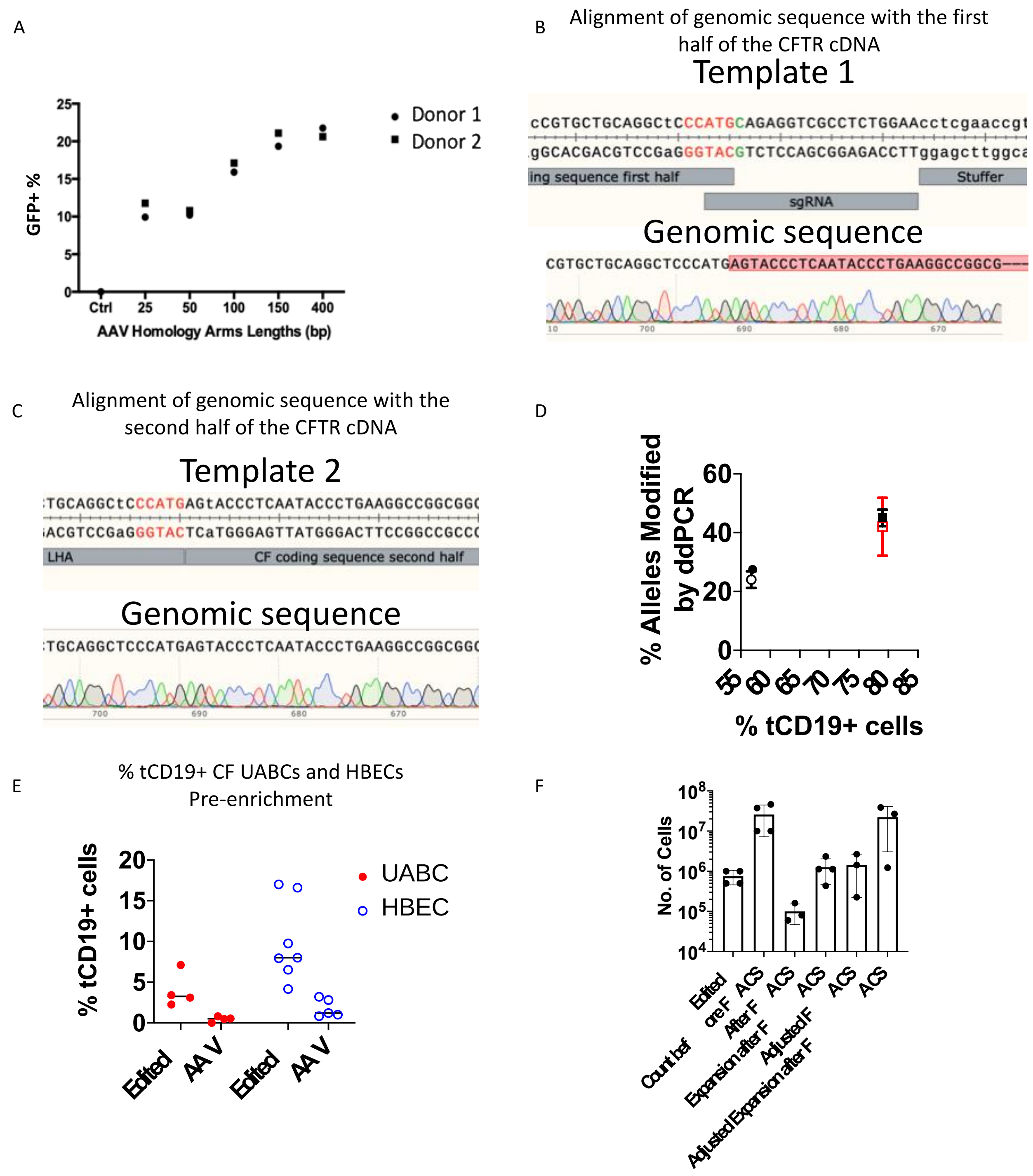


**Fig. S1.** Characterization of edited and enriched UABCs. (**A**) HR templates of different homology arm lengths coding for GFP were inserted into the CFTR locus of UABCs. Percentage of GFP$^+$ UABCs was ~10% for HAs under 100 bp and reached ~20% for 150 bp HA and did not increase further with a longer HA of 400 bp. (**B**) The junction spanning the two HR templates corresponding to the two halves of the CFTR cDNA was amplified using PCR and analyzed by Sanger sequencing. The genomic sequence shows the presence of the sequence right up to the end of the first half of the CFTR cDNA highlighted in red and the absence of the sgRNA sequence and the stuffer sequence present in the HR template. (**C**) Genomic sequence from the same donor shows the seamless integration of the second half of the CFTR cDNA with the first half. (**D**) Allelic correction rates of the corrected UABCs were compared with the percent of cells that were $tCD19^+$. Each symbol represents cells from a separate donor. The error bars represent the technical variation observed in duplicate runs of the ddPCR assay. The percentage of modified alleles is ~50% of the percentage of $tCD19^+$ cells suggesting monoallelic integration of the CFTR cDNA. (**E**) UABCs and HBECs from donors with CF were edited to insert the CFTR cDNA and the tCD19 expression cassette. 4 $\pm$ 2% UABCs were $tCD19^+$ and 10 $\pm$ 5% HBECs were $tCD19^+$. (**F**) Number of UBACs at each step in the correction process. Starting from 0.5-1 million cells, UABCs were edited and expanded to obtain 10-40 million for FACS enrichment. Only 3-5 million cells out of the 10-40 million cells were enriched using FACS and we obtained ~1 million UABCs one passage after FACS enrichment. The estimated yield that could have been obtained if all the edited cells were enriched using FACS and then expanded are also provided. These columns are labeled as adjusted FACS and adjusted expansion after FACS.

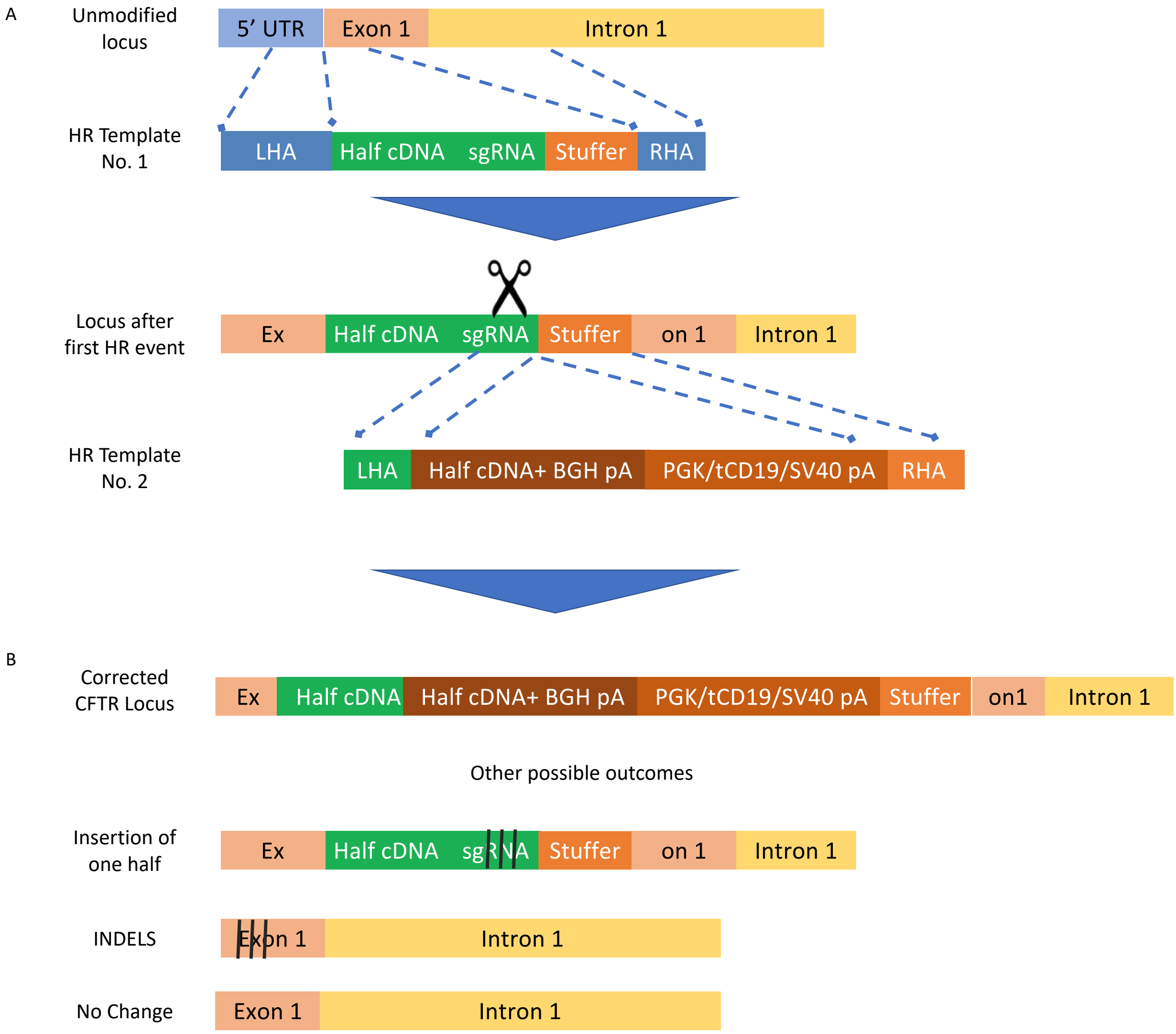


**Fig. S2.** Sequential insertion of the CFTR cDNA by two homologous recombination events. (**A**) The Cas9 RNP/sgRNA complex first induces a break in exon 1. The first HR template contains left and right homology arms (HA) of 400 bp length that resemble the 5’ UTR region of the CFTR locus and part of exon 1 and intron 1 respectively. The first HR template also contains a stuffer sequence that will act as a HA for the second HR event. After the first HR event, a modified locus will consist of the first half of the CFTR cDNA and stuffer sequence. The second HR template contains the second half of the CFTR cDNA followed by a BGH poly A tail and a tCD19 expression cassette consisting of the PGK promoter, tCD19 and SV40 polyA tail. The LHA and RHA consist of the last 400 bp of the first HR template and the stuffer respectively. (**B**) A fully corrected allele will contain the full CFTR cDNA, a BGH polyA , followed by a PGK promoter, truncated CD19 and an SV40 poly A tail. It is likely most of the enriched cells contain this modification in only one allele. The other allele is likely to contain other possible outcomes including the first half of the insert (with or without INDELs in the end), INDELs in the beginning of Exon 1 (no HR) or an unmodified allele. The sgRNA was screened to induce INDELs in over 90% of alleles in the absence of an HR template. It is therefore likely that the presence of an unmodified locus in the second allele is extremely rare. INDELs are indicated by three black lines in the figure.

A

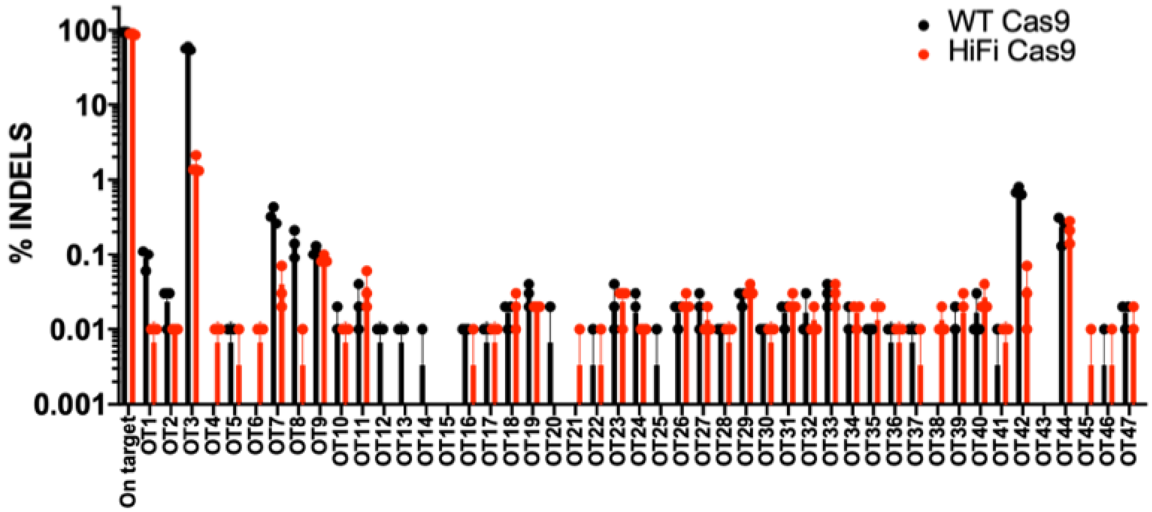


**Fig. S3.** Off-target activity of the sgRNA. (**A**) The sgRNA targeting the exon 1 of CFTR locus shows OT activity in one locus (OT-3) when airway basal stem cells are edited using wild-type (WT) Cas9. The use of high-fidelity (HiFi) Cas9 reduced the OT activity significantly in OT-3.

The percent of alleles with INDELs was reduced from ~50% when using WT-Cas9 to ~1% when using HiFi Cas9 while the percent INDELs in the on-target site remained the same (>85%).
